# Development of an orthotopic medulloblastoma zebrafish model for rapid drug testing

**DOI:** 10.1101/2024.02.21.578208

**Authors:** Niek van Bree, Ann-Sophie Oppelt, Susanne Lindström, Leilei Zhou, Lola Boutin, John Inge Johnsen, Lars Bräutigam, Margareta Wilhelm

## Abstract

Medulloblastoma (MB) is one of the most common malignant brain tumors in children. Current preclinical *in vivo* model systems for MB have increased our understanding of molecular mechanisms regulating MB development; however, they may not be suitable for high-throughput screening efforts. We demonstrate here that transplantation of seven different MB cell lines or patient-derived cells into the blastula stage of zebrafish embryos leads to orthotopic tumor cell growth that can be observed within 24 hours after transplantation. Importantly, the homing of transplanted cells to the hindbrain region and the aggressiveness of tumor growth are enhanced by pre-culturing cells in a neural stem cell-like medium. The change in culture conditions rewires the transcriptome towards a more migratory and neuronal progenitor phenotype, including the expression of guidance molecules SEMA3A and EFNB1, both of which correlate with lower overall survival in MB patients. Furthermore, we highlight that the orthotopic zebrafish MB xenograft model has the potential to be used for high-throughput drug screening.

**Key points:**

1. Medulloblastoma cells home to the hindbrain region in developing zebrafish embryos.
2. Neural stem cell culture conditions improve the homing capacity of MB tumor cells.
3. Medulloblastoma-transplanted zebrafish embryos can be used as a high-throughput *in vivo* model for drug screening.

**Importance of the Study:** One of the challenges of accurately modeling medulloblastoma is the large heterogeneity in tumor characteristics. To accurately model this heterogeneous disease, patient-derived xenograft mouse models are currently the standard. However, such mouse models are labor intensive, time-consuming, and not suitable for high-throughput studies. Here, we describe a quick and straightforward zebrafish xenograft model that provides a promising alternative to these existing mouse models. We demonstrate that this model can be utilized to study tumor cell growth of several major medulloblastoma subgroups. More importantly, our model facilitates high-throughput drug testing, providing a scalable opportunity for *in vivo* drug screenings that will support the discovery of novel therapeutic compounds against medulloblastoma.

## Introduction

Medulloblastoma (MB) is one of the most common malignant pediatric brain tumors. Advances in molecular characterization and innovations in the multimodal treatment of MB have drastically increased the survival rate, and today about 70% of patients are cured. However, prognosis is strongly correlated with the molecular MB subgroups (Wingless (WNT), Sonic Hedgehog (SHH), Group 3, and Group 4), with patients with Group 3 MB having the worst prognosis [1]. This suggests that more subgroup-specific treatments are needed. Furthermore, the improvement in survival has stalled and remained relatively stable over the past two decades [2]. In addition, approximately 30% of MB patients (most commonly belonging to the SHH, Group 3, or Group 4 subgroup) still relapse, which is almost always fatal [3]. To further improve the survival rate of MB patients and identify novel treatment options, new disease-specific models are required. Current *in vivo* models for MB include genetically engineered mouse models, orthotopic transplants, and patient-derived xenograft mouse models [4]. While these mouse models accurately represent MB subgroups, they are labor intensive, time-consuming, and subject to ethical and financial constraints that make them impractical for use in large-scale studies.

Recently, zebrafish (*Danio rerio*) embryos have emerged as a promising *in vivo* model for many human cancers, including non-small cell lung cancer brain metastasis, pilocytic astrocytoma, and glioblastoma [5-7]. Zebrafish have proven to be a good model system for high-throughput transplantation studies because of their high fecundity, *ex utero* embryogenesis, small size, rapid development, and low maintenance and husbandry costs [8]. Furthermore, previous studies have shown that cell line-derived (CDXs) and patient-derived xenografts (PDXs) in zebrafish embryos are able to proliferate *in vivo* and display disease and patient-specific characteristics, suggesting that zebrafish models can be used as an alternative to existing mouse models and could enable rapid identification of patient-specific targeted treatments [7, 9-13]. These favorable pre-clinical results have led to the initiation of several trials to study the translational and clinical use of zebrafish PDXs as a model for tumor growth and treatment efficacy (NCT03668418, NCT05289076, NCT05616533). However, despite all promising results of zebrafish cancer xenografts, orthotopic zebrafish models have not been explored for MB.

Here, we report the use of zebrafish embryos as an orthotopic model for MB. We demonstrate that MB CDXs and PDXs home primarily to the hindbrain region after transplantation into the blastula-stage of developing zebrafish embryos, independently of MB subgroup. This homing mechanism can be further enhanced by conditioning cells to neural stem cell conditions containing growth factors and supplements important for neuronal growth and synaptic plasticity. Furthermore, we show that MB-transplanted zebrafish embryos have the potential to be used as a high-throughput orthotopic MB zebrafish model for testing drug compounds.

## Materials and Methods

### Animal models

8-to 12-week-old female NOD/SCID/IL2Rγ^-/-^ (NSG) mice were used for orthotopic transplantation of 1×10^5^ secondary tumor NES (tNES) cells and tumor growth was monitored using IVIS SpectrumCT In Vivo Imaging System (PerkinElmer) as previously described [14]. Mice were euthanized upon observation of ethical endpoint criteria. fli1:EGFP zebrafish embryos aged about 4 hours old (1k-cell stage) were used for transplantation. Fertilized eggs were obtained from natural spawning and staged according to standard procedures [15]. Transplanted zebrafish embryos were kept for a maximum of 3 days after transplantation. Experiments on embryos younger than 5 days do not require an ethical permit. All zebrafish experiments were conducted in accordance with the national ethical guidelines and regulations (N207/14). Mouse experiments were approved by Stockholm’s North Ethical Committee of Animal Research (ethical permit 6548/18).

### Generation of tertiary tumor NES cell lines

Tertiary tNES cells were isolated from established tumors from orthotopic transplantation of secondary tNES cells (Supplementary Figure 1). Tumor cell isolation was performed as previously described [14]. In brief, tumor tissue was dissociated using digestion buffer (1x PBS with 10 U/ml papain, 200 μg/ml L-cysteine, 250 U/ml DNAse, 1% penicillin-streptomycin, all from Sigma). Cells were triturated and passed through a 100 μm cell strainer to obtain a single-cell suspension. Mouse cell contaminations were removed using the Mouse Cell Depletion Kit (Miltenyi Biotec, 130-104-694). Tertiary tNES cells were cultured in complete neural stem cell medium (see section Cell culture) on poly-L-ornithine/laminin-coated plates.

**Figure 1.**
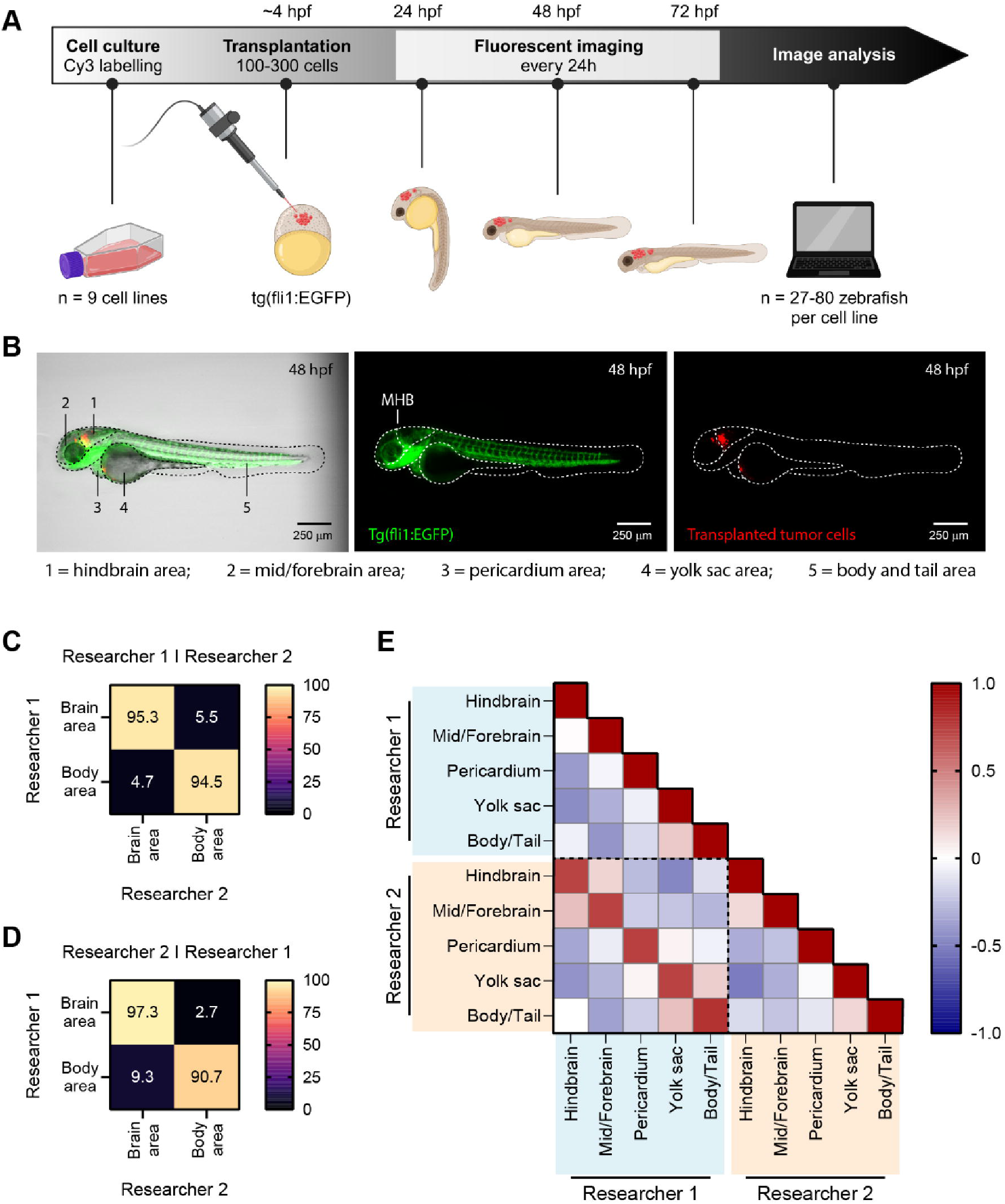
Homing of fluorescently labeled tumor cells transplanted in zebrafish embryos can be examined with fluorescent imaging and image analysis. **A)** Schematic overview of the experimental workflow for zebrafish embryo transplantations followed by tumor location analysis. **B)** Representative image of a tertiary tNES cell transplanted zebrafish embryo 48 hours post fertilization (hpf), displaying the five defined locations for location analysis. Left panel: bright field with GFP and Cy3 overlay. Middle panel: expression of enhanced GFP in the vasculature of the fli1:EGFP zebrafish embryo (GFP, green). MHB = mid-hindbrain boundary. Right panel: labeled tumor cells (Cy3, red). **C)** Confusion matrices of image classification performed by researcher 1, **D)** and researcher 2. **E)** Pearson correlation matrix between the analyzed datasets of researcher 1 and researcher 2 revealing high correlation between individually analyzed datasets.

### Cell culture

Tertiary tumor-isolated neuroepithelial stem (tNES) cells were cultured as previously described [14]. Briefly, tNES cells were cultured on flasks coated with 20 μg/ml poly-L-ornithine (Sigma, P3655) and 1 μg/ml laminin (Sigma, L2020) in complete neural stem cell medium (DMEM/F12+Glutamax (ThermoFisher, 31331093) supplemented with 1% N2 supplement (ThermoFisher, 17502001), 0.1% B27 supplement (ThermoFisher, 17504044), 10 ng/ml FGF2 (Qkine, Qk053), 10 ng/ml EGF (PeproTech, AF100-15) and 1% penicillin-streptomycin (Sigma, P4333)). The PDX cell line MB-LU-181 and CHLA-01-MED cells were cultured in DMEM/F12+Glutamax with 20 ng/ml (CHLA-01-MED) or 40 ng/ml (MB-LU-181) FGF2, 20 ng/ml EGF, 2% B27, and 1% penicillin-streptomycin. DAOY, ONS-76, MDA-MB-231, and HCT116 p53^+/+^ cells were maintained in DMEM (ThermoFisher, 41966052) supplemented with 10% FBS (ThermoFisher, SV30160.03HI) and 1% penicillin-streptomycin.

Additionally, medium for DAOY cells contained 1% MEM non-essential amino acids, 1% HEPES, and 1% Glutamax (ThermoFisher, cat. 11140050, SH30237.01, and 35050061, respectively). UW228-3 cells were cultured in RPMI (ThermoFisher, 61870044) and D425 cells in DMEM/F12+Glutamax, both supplemented with 10% FBS and 1% penicillin-streptomycin. All cell lines tested negative for mycoplasma and were authenticated by STR analysis at Eurofins (DAOY, D425, CHLA-01-MED), ECACC (MDA-MB-231, HCT116 p53^+/+^), or Multiplexion (ONS-76, UW228-3). MB-LU-181 PDX establishment has previously been described in [16].

### Lentiviral transduction

Luciferase expressing cells were generated through lentiviral transduction of pLenti-CMV-Luc-Puro. Lentiviral particles were produced in HEK293T cells using packaging and envelope constructs pCMVΔ8.2 and pMD.G-VSV-G (pCMVΔ8.2 and pMD.G-VSV-G were gifts from Bob Weinberg, Addgene plasmids #8454, #8455), and concentrated using fast-trap virus purification and concentration kit according to manufacturer′s instructions (Millipore). For transduction, MB cells were seeded into 6-well plates and transduced with virus at 70% confluency, 24h after of transduction, cells were selected with 2 μg/mL puromycin.

### Cell proliferation assay

To assess cell proliferation over time, UW228-3 cells were seeded in 12-well tissue culture treated plates at a density of 5×10^4^ cells/well. MDA-MB-231 and D425 cells were seeded at a density of 1×10^5^ cells/well. Cells were dissociated using TryplE Select (ThermoFisher, A1217701) and counted at defined time points to determine the number of viable cells.

### Transplantation of cell cultures and treatments

Confluent cell cultures were labeled with Vybrant™ DiI cell-labeling solution (ThermoFisher, V22885) in the corresponding culture medium (1:200) for 15-20 minutes according to manufacturer’s instructions. Cells were washed with PBS for 5 minutes, followed by trypsinization and centrifugation at 300g for 3 minutes. Cells were resuspended and centrifuged twice to assure complete removal of the dye. Fluorescent labeled tumor cells were transplanted in the blastula-stage of fli1:EGFP zebrafish embryos as previously described [17]. In brief, embryos were lined up on a 1% agarose mold in E3 medium. Directly before transplantation, cell suspensions were spun down and almost all medium was removed. Concentrated cell suspension was loaded into a microcapillary (World Precision Instruments, TW100-4) connected to a FemtoJet 4x (Eppendorf). About 100-300 cells were injected in the center of the cell mass. After transplantation, the embryos were collected in E3 medium and raised at 33°C. Embryos with intracranial tumors were selected for treatment 24 hours aftertransplantation. LDE225 phosphate (sonidegib, Cayman Chemicals, 16263) and 4-hydroxycyclophosphamide (4-HCP, Santa Cruz Biotechnology, sc-206885A) were added directly in E3 medium to a final concentration of 10 μM. DMSO (Sigma, D8418) was used as a control.

### High-throughput imaging of zebrafish embryos

For high-throughput imaging of the transplanted zebrafish embryos, imaging 96-well plates (Ibidi, 89621) with agarose molds were prepared [18]. Single embryos were distributed into the wells together with 150 μl of exposure medium (160 μg/ml tricaine, 30 mg/L phenylthiourea in E3 medium). Embryos were manually oriented into position and imaged using ImageXpress Nano (Molecular Devices) every 24 hours. After imaging, the plate was moved into a wet chamber inside an incubator at 33°C. Zebrafish embryo images were analyzed for tumor location (hindbrain, mid/forebrain, pericardium, yolk sac, and body/tail area) using ImageJ 1.53. Unfocused images or dead embryos were excluded from the analysis. All data was analyzed by two people individually of which researcher 2 was blinded. To compare the two analyzed datasets, each image has been quantitatively classified from area with the highest percentage of tumor cells (=1) to area with the lowest percentage of tumor cells (=5).

### Luciferase activity measurement

To measure luciferase activity, treated embryos were transferred into a white-bottom 96-well plate (Greiner, 655074). Exposure medium was removed before adding 100 μl of freshly prepared lysis buffer containing 10% glycerol, 1% Triton-X 100, 25 mM Tris-phosphate pH 7.8 adjusted with phosphoric acid, 8 mM magnesium chloride, and 1mM DL-dithiothreitol (Sigma, cat# G012, T8787, T1503, 438081, M8266, 43815, respectively) in dH_2_O. The plate was placed on a rocking platform and embryos were lysed for 45 minutes at room temperature. 100 μl of freshly prepared substrate buffer, 25mM Tris-phosphate pH 7.8, 1mM DL-dithiothreitol, 0.25 mg/ml D-luciferin (BioThema, BT11-1000K), and 1 mM ATP (ThermoFisher, R0441) in dH_2_O, was added and incubated for 5 minutes in the dark at room temperature. Luminescence was measured using the FLUOstar Omega (BMG Labtech).

### Transcriptome sequencing

UW228-3 and D425 cells were gradually conditioned towards complete neural stem cell medium by increasing the concentration of this medium from 25% to 100% over a course of four passages. Neural stem cell medium for UW228-3 cells was supplemented with 1% FBS to prevent cell death. Conditioned UW228-3 and D425 cells were cultured for at least two passages in complete neural stem cell medium before isolating total RNA for RNAseq. Total RNA was extracted using TRIzol Reagent (ThermoFisher, 15596018) and Direct-zol RNA extraction kit (Zymo Research, R2052) according to manufacturer’s instructions. RNA sequencing was performed using the Illumina Novaseq 6000 (PE150) platform. RNA sequence reads were mapped against the human reference genome GRCh38/hg38 using Hisat2 v2.0.5 [19]. Differential expression analysis of two groups (three biological replicates per group) was performed using the DESeq2 R package (1.20.0). DESeq2 provides statistical routines for determining differential expression in digital gene expression data using a model based on the negative binomial distribution. The resulting P-values were adjusted using the Benjamini and Hochberg’s approach for controlling the false discovery rate. Genes with an adjusted P-value <=0.05 found by DESeq2 were assigned as differentially expressed.

### Gene set enrichment analysis

Gene Set Enrichment Analysis (GSEA) was performed using GSEA v.4.2.3 [20]. The transcriptome sequencing dataset was analyzed against the MSigDB Hallmark, BioCarta, KEGG, Pathway Interaction Database (PID), Reactome, WikiPathways, and Gene Ontology (GO) gene sets according to Reimand et al. [21]. Pathways enrichment analysis and comparison of multiple transcriptome datasets was performed by using Metascape [22].

### Statistical analysis

Data is presented as means ± standard deviation (SD) or standard error of the mean (SEM). Statistical significance was analyzed with the Student *t* test, one-way or two-way ANOVA with Dunnett, Tukey, or Šídák post-hoc test. P-values <0.05 were considered statistically significant. All statistical analyses were performed using GraphPad Prism 9.5.0. *P < 0.05, **P < 0.01, ***P < 0.001, ****P < 0.0001.

## Results

### Tumor cell transplantations into the blastula-stage of zebrafish embryos as an in vivo model for medulloblastoma

Previous studies have observed that human melanoma and glioblastoma cells are able to home to their environment of origin [17, 23]. It was proposed that intrinsic morphogenetic cues in the early developing zebrafish embryo are responsible for this homing mechanism by providing support to these human cells. Based on these results, we speculated that MB cells can migrate to the developing cerebellum after transplantation into the blastula-stage of zebrafish embryos (Figure 1A). We transplanted a total of seven fluorescently labeled MB cell lines (both PDXs and established cell lines) and two non-MB cell lines into fli1:EGFP embryos four hours post fertilization (hpf). Transgenic fli1:EGFP zebrafish express GFP in vascular endothelial cells. The transplanted zebrafish embryos were imaged every 24 hours for three days after fertilization to track tumor cell localization. Remarkably, the majority of tumor cells were already present in the region of interest 24 hpf. To quantify tumor localization, images acquired 48 hpf were used in which the developing embryo was divided into five areas, namely hindbrain, mid/forebrain, pericardium, yolk sac, and body/tail area (Figure 1B, Supplementary Figure 2A). One of the most distinct veins in the early developing zebrafish embryo is the mid-cerebral vein, which delineates the mid-hindbrain boundary (MHB) [24], and helps us to separate the hindbrain area from the mid/forebrain area without the need for closeup images of the head.

**Figure 2.**
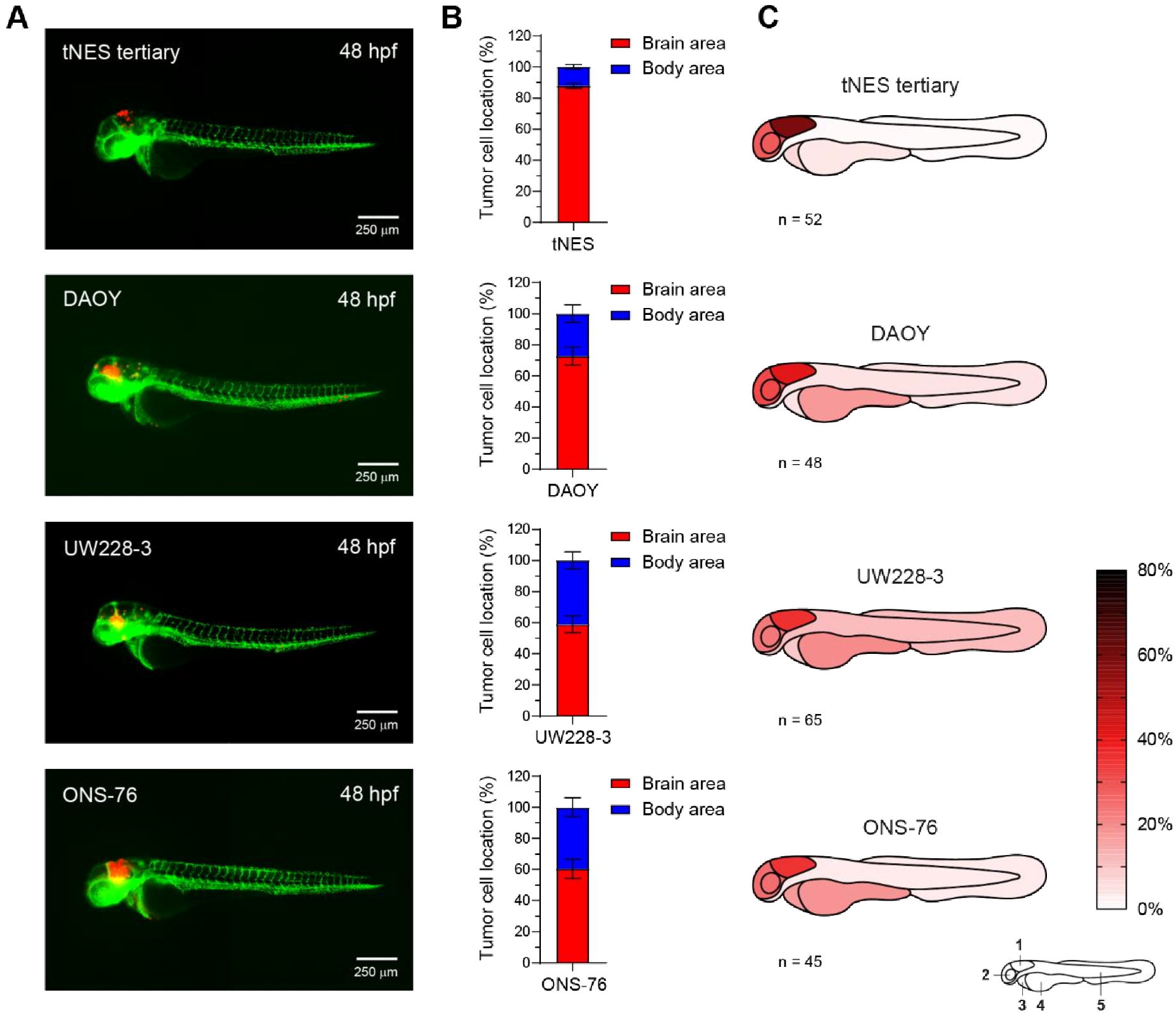
SHH medulloblastoma cell lines home towards the hindbrain region after transplantation into the blastula-stage of zebrafish embryos. **A)** Representative images of fli1:EGFP zebrafish embryos 48 hpf transplanted with SHH medulloblastoma (MB) cell lines (tertiary tNES, DAOY, UW228-3, and ONS-76). **B)** Brain/body area distribution of transplanted SHH MB cell lines in 48 hours old embryos. Mean ± SEM. **C)** Schematic heatmaps of the tumor cell location of SHH MB cell lines in 48 hours old embryos. 1 = hindbrain area; 2 = mid/forebrain area; 3 = pericardium area; 4 = yolk sac area; 5 = body and tail area.

To ensure that the analysis of tumor localization was performed accurately and without bias, all transplanted embryos (*n =* 666) were analyzed by two researchers, the second researcher of whom was blinded to cell line identity. Classification by defining majority of the tumor cells in either the brain or body area resulted in a high agreement between both researchers (95.0%) (Figure 1C and 1D), and good accuracy (77.2%) was achieved even with deeper classification into five anatomical areas (Supplementary Figure 2B and 2C). When examining the differently classified images, no major differences in the percentiles of tumor localization were observed (Supplementary Figure 2D), and even embryos with a high spread of tumor localization were analyzed uniformly (Supplementary Figure 2E). Further analysis using Pearson correlation revealed that all five anatomical areas showed a strong positive correlation when comparing the two analyzed datasets (Figure 1E). As both analyses were very similar, average values were used for further presentation of the data.

### Medulloblastoma tumor cells home towards the hindbrain region in developing zebrafish embryos

First, we transplanted tertiary tumor neuroepithelial stem (tNES) cells into the blastula-stage of zebrafish embryos to test our hypothesis. Tertiary tNES cells represents malignant human SHH-MB and are derived from a stem cell model that we previously established by reprogramming non-cancerous skin cells carrying a germline *PTCH1* mutation and differentiating them into patient-derived NES cells [14]. These patient-derived NES cells form MB tumors upon orthotopic transplantation into the cerebellum of NSG mice from which tNES cells were isolated, serial re-transplantation generates more malignant tNES (Supplementary Figure 1). Interestingly, we observed a high degree of brain localization when tertiary tNES were transplanted, with 87.9 ± 1.6% homing to the brain region (Figure 2A and 2B). Further analysis revealed that 67.3% of the tertiary tNES cells that were found in the brain were localized in the hindbrain region (Figure 2C). To determine whether this ability to localize to the hindbrain is a universal property of SHH-MB cells, we transplanted DAOY, UW228-3, and ONS-76 cells in blastula-stage embryos. Like tertiary tNES cells, we observed that the majority of tumor cells were located to the brain for all three SHH-MB cell lines at 48 hpt: 72.8 ± 5.7% of DAOY cells, 59.1 ± 5.4% of UW228-3 cells, and 60.5 ± 6.2% of ONS-76 cells (Figure 2A and 2B).

Again, a considerable proportion of these brain-localized tumor cells localized to the hindbrain region (57.0%, 60.6%, and 58.5% for DAOY, UW228-3, and ONS-76 cells, respectively) (Figure 2C). After observing the high degree of hindbrain localization of SHH-MB tumor cells, we speculated whether this is a global MB mechanism. To investigate the localization of other MB subgroups, we transplanted the Group 3 MB PDX cell line MB-LU-181 [16]. Importantly, transplantation of these cells also resulted in a high percentage of tumor cells located in the brain area with 81.5 ± 6.1% (*n =* 27) (Figure 3A and 3B). The area-specific evaluation showed that 75.0 % of these cells were localized in the hindbrain area (Figure 3C). Additional transplantation with the Group 3 MB cell line D425 and the Group 4 MB neurosphere cell line CHLA-01-MED [25] yielded comparable results and showed a high proportion of tumor cells localized in the brain area, 62.0 ± 3.7% (*n =* 80) for D425 and 86.4 ± 2.9% (*n =* 70) for CHLA-01-MED, with a high proportion of these cells confined within the hindbrain area (64.5% for D425 and 83.8% for CHLA-01-MED) (Figure 3A, 3B, and 3C). To study if the homing mechanism is specific to MB tumor cells or if it is a common process among other cancers, we transplanted the breast cancer cell line MDA-MB-231 and the colon cancer cell line HCT116 p53^+/+^. Both non-MB cell lines had significantly lower levels of brain localization compared to the MB cell lines, with only 29.1 ± 4.4% (*n =* 67) of MDA-MB-231 and 32.8 ± 4.6% (*n =* 72) of HCT116 p53^+/+^ cells detected in the brain area (Figure 3D and 3E and Supplementary Figure 3B). Instead, most non-MB cancer cells localized to the non-brain area with preference to the yolk sac, with 43.3 ± 4.4% for MDA-MB-231 and 31.3 ± 4.4% for HCT116 p53^+/+^ (Figure 3F). Intriguingly, different types of embryonal malformations were more commonly observed upon transplantations with non-MB cell lines compared to MB cell lines. Zebrafish embryos transplanted with MB cell lines displayed cranial malformations more often whereas pericardial edema and yolk sac edema were more frequently detected in zebrafish embryos transplanted with non-MB cells (Supplementary Figure 3A). Taken together, these results suggest an intrinsic homing mechanism of MB tumor cells upon transplantation in developing zebrafish embryos.

**Figure 3.**
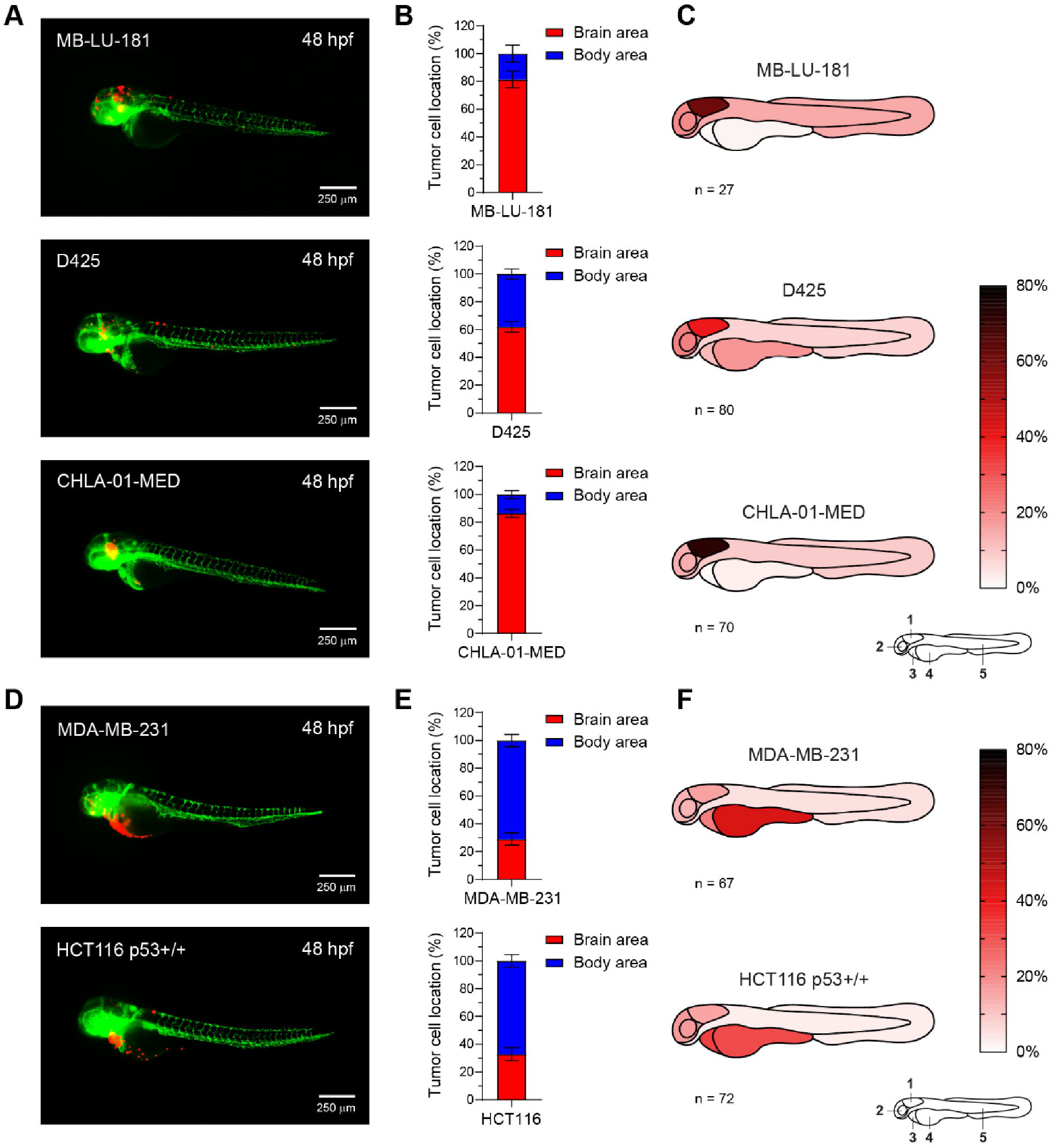
Group 3 and Group 4 medulloblastoma cell lines display homing towards the hindbrain area upon transplantation into the blastula-stage of zebrafish embryos whereas non-medulloblastoma cell lines do not. **A)** Representative images of fli1:EGFP zebrafish embryos 48 hpf transplanted with Group 3 (MB-LU-181 and D425), Group 4 (CHLA-01-MED) MB, **D)** and non-MB cell lines (MDA-MB-231 and HCT116 p53^+/+^. **B)** Brain/body area distribution of transplanted Group 3 and Group 4 MB cell lines and **E)** non-MB cell lines in 48 hours old embryos. Mean ± SEM. **C)** Schematic heatmaps of the tumor cell location of Group 3 and Group 4 MB, **F)** and non-MB cell lines in 48 hours old embryos. 1 = hindbrain area; 2 = mid/forebrain area; 3 = pericardium area; 4 = yolk sac area; 5 = body and tail area.

### Neural stem cell culture conditions improve homing capacity of tumor cells

Interestingly, we found that the patient-derived MB cell lines (tNES and MB-LU-181) and the neurosphere cell line CHLA-01-MED had a significantly higher homing capacity towards the brain compared to the other transplanted MB cell lines (DAOY, UW228-3, ONS-76, and D425) (Supplementary Figure 3B). Since all three of these cell lines are cultured in neural stem cell condition, we adapted one SHH-MB cell line (UW228-3) and one Grp3 cell line (D425) to similar growth conditions prior to transplantation (Supplementary Figure 4A). Interestingly, MB cells adapted to neural stem cell media (UW228-3+ and D425+) showed significantly higher localization to the brain area and specifically to the hindbrain area compared to cells cultured in standard cell culture conditions (Figure 4A, 4B, and 4C). This was specific to MB cells, since MDA-MB-231 breast cancer cells adapted to neural stem cell conditions did not show any significant increase in homing to the brain area. Instead, we observed a significant increase of cell localizing to the pericardium area (Figure 4A, 4B, and 4C). Switching the cells to neural stem cell conditions resulted in a decrease in cell proliferation *in vitro* for UW228-3+ and MDA-MB231+ cells. However, it caused a more aggressive disease progression *in vivo* in transplanted zebrafish embryos (Figure 4D and 4E), resembling the disease progression of the patient-derived MB cell lines (tNES and MB-LU-181) (Supplementary Figure 4B and 4C). These observations suggest that growth conditions play an instrumental role in providing correct homing cues.

**Figure 4.**
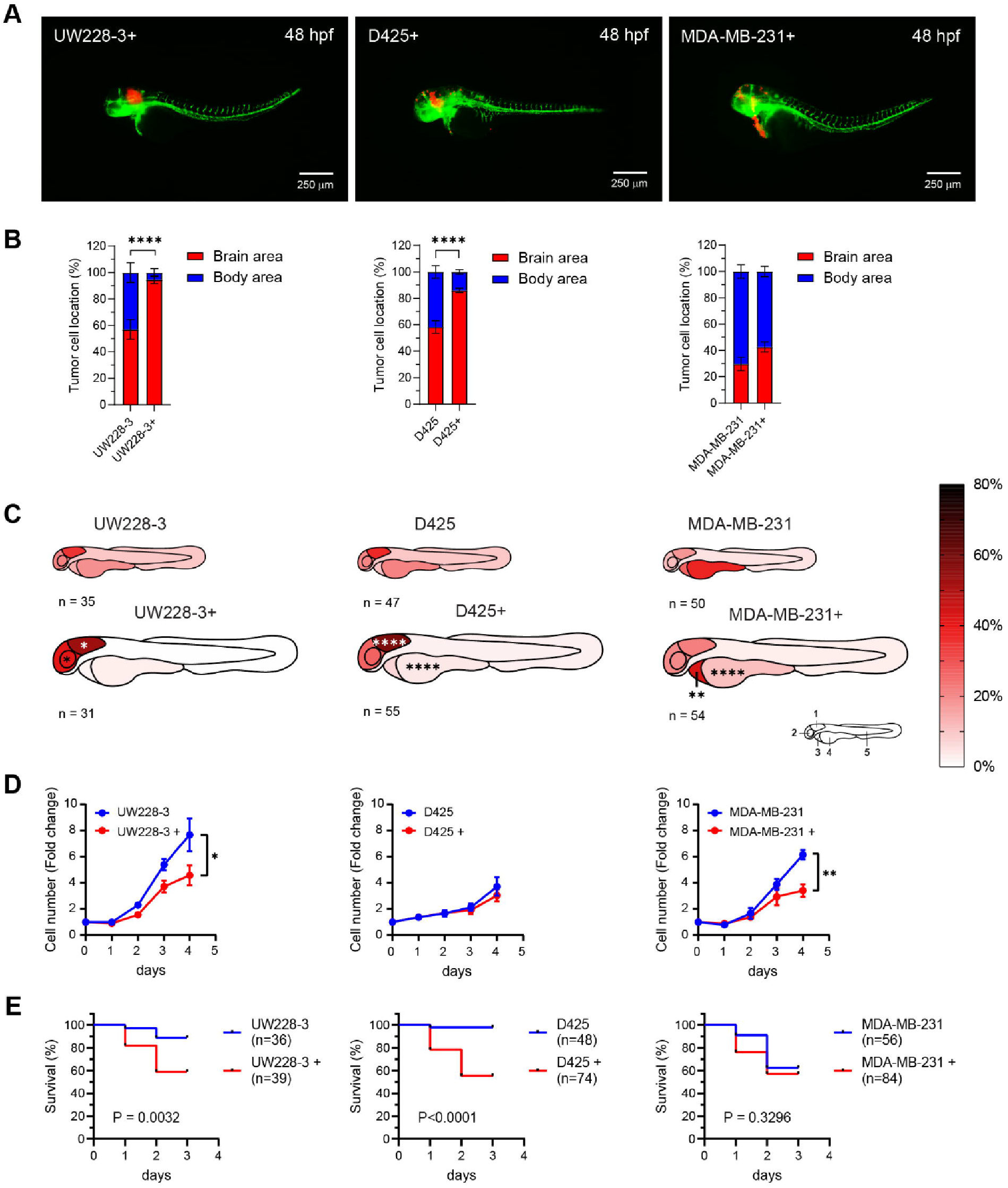
Culture medium composition has a significant effect on tumor cell localization in transplanted zebrafish embryos. **A)** Representative images of fli1:EGFP zebrafish embryos 48 hpf transplanted with tumor cell lines cultured in complete neural stem cell medium (UW228-3+, D425+, and MDA-MB-231+). Medium for UW228-3+ was supplemented with 1% FBS for maintenance and removed before transplantation. **B)** Brain/body area distribution of tumor cell lines cultured in complete neural stem cell medium (+) compared to normal culture medium in 48 hours old embryos. Mean ± SEM, ****P<0.0001, Student *t* test. **C)** Schematic heatmaps of the tumor cell location of tumor cell lines cultured in complete neural stem cell medium (+) compared to normal culture medium in 48 hours old embryos. 1 = hindbrain area; 2 = mid/forebrain area; 3 = pericardium area; 4 = yolk sac area; 5 = body and tail area. *P<0.05, **P<0.01, ****P<0.0001, two-way ANOVA with Šídák post-hoc test. **D)** Proliferation rate of UW228-3, D425, and MDA-MB-231 cells cultured in medium with FBS or in neural stem cell medium (+) evaluated by cell counting. Mean ± SD, n = 3 independent experiments. **E)** Overall survival analysis of zebrafish transplanted with UW228-3, D425, and MDA-MB-231 cells cultured in medium with FBS or in neural stem cell medium (+). Kaplan-Meier curves depict differences in survival and statistical differences were determined using the Log-rank Mantel-Cox test.

To gain a better understanding of what confers increased homing capacity in cells cultured in neural stem cell conditions, we analyzed transcriptional changes in UW228-3 and D425 cultured under standard culture conditions compared with neural stem cell culture conditions (Figure 5A). Differential gene expression identified 1289 genes with significantly altered expression (616 genes upregulated, 673 downregulated, FDR<0.05, FC>±2) in UW228-3+ cells compared to UW228-3 (Figure 5B, Supplementary Dataset 1), and 984 genes (667 genes upregulated, 317 downregulated, FDR<0.05, FC>±2) in D425+ cells compared to D425 (Figure 5C, Supplementary Dataset 1).

**Figure 5.**
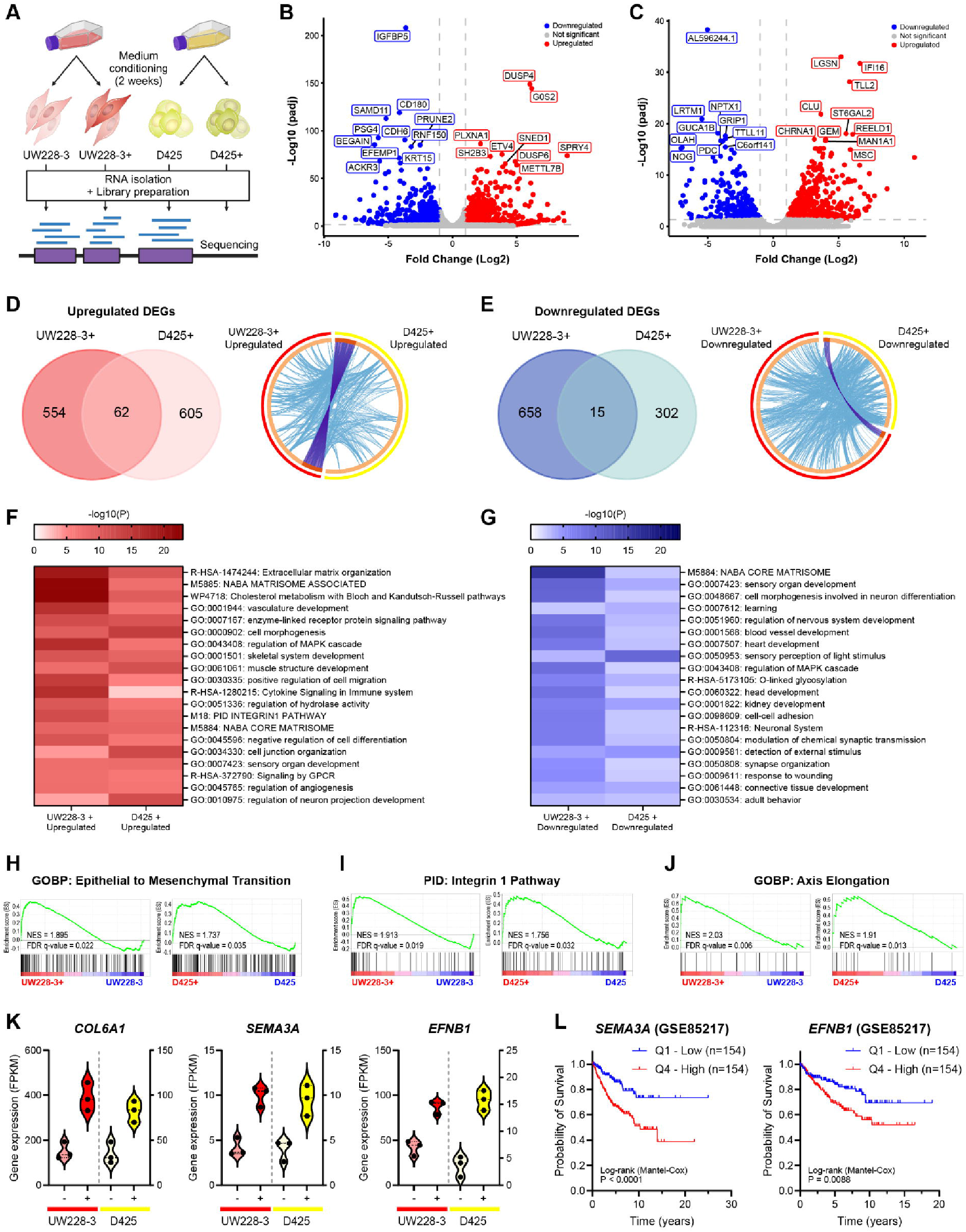
Transcriptomic analysis reveals a myriad of upregulated gene pathways associated with cancer stemness and migration. **A)** Schematic overview of the bulk RNA sequencing protocol. **B)** Volcano plot showing all differentially expressed genes (DEGs) upon comparison of the transcriptome of UW228-3 with UW228-3 in complete neural stem cell medium (+), or **C)** D425 with D425 cultured in complete neural stem cell medium (+). Gene names of the top 20 most significant DEGs are shown. Significant DEGs are those with a fold change of ≥2 (Log2FC≥1) and an adjusted P-value of ≤0.05. Blue dots represent downregulated genes and red dots represent upregulated genes. **D)** Left panel: Venn diagram showing overlap between the upregulated and **E)** downregulated DEGs in UW228-3+ and D425+. Right panel: Circos plots displaying the gene overlap and shared term level between the gene lists UW228-3 versus UW228-3+ and D425 versus D425+. Purple lines link identical genes and blue lines link genes that belong to the same enriched ontology term. The inner circle represents the two gene lists, whereas the outside circle represents the group. **F, G)** Heatmaps of enriched gene set terms shared across both gene lists, colored by P-values. Each term represents a cluster of multiple associated gene sets. Clusters were named after the most significantly expressed gene set present in both gene lists. **H-J)** Epithelial to mesenchymal transition, integrin 1 pathway, and axis elongation are all significantly enriched in both UW228-3 and D425 when cultured in complete neural stem cell medium (+) as observed upon GSEA analysis. **K)** Violin plots showing the DEGs *COL6A1, SEMA3A*, and *EFNB1* that are significantly enriched in both UW228-3+ and D425+, expressed as fragments per kilobase (FPKM). **L)** Kaplan Meier curves represent the survival of MB patients with low expression (Q1) compared with MB patients with high expression (Q4) of *SEMA3A* or *EFNB1*.

Surprisingly, there was little overlap between the differentially regulated genes in UW228-3+ and D425+ cells, with only 62 upregulated genes and 15 downregulated genes shared between them (Figure 5D and 5E, left panels). However, pathway enrichment analysis identified that UW228-3+ and D425+ share a high degree of genes belonging to the same enriched ontology term (Figure 5D and 5E, right panels) and converge on a multitude of similar gene sets (Figure 5F and 5G). Importantly, many of the upregulated pathways were associated with enhanced migratory capacity, such as pathways involved in extracellular matrix organization, matrisome pathways, integrin pathway, and the positive regulation of cell migration (Figure 5F, Supplementary Figure 5A, Supplementary Dataset 2). Other upregulated gene set terms were associated with organotypic developmental programs, including pathways related to negative regulation of cell differentiation and regulation of neuron projection development. This suggests that established MB cell lines can be reconditioned to a more progenitor-like state. Additionally, downregulation of pathways involved in the regulation of nervous system development, neuron differentiation, and cell-matrix or cell-cell adhesion may further increase the migratory capacity and promote the progenitor-like state of UW228-3+ and D425+ (Figure 5G, Supplementary Figure 5B, Supplementary Dataset 3). This was further validated using gene set enrichment analysis (GSEA), which showed that among others epithelial to mesenchymal transition and the integrin-1 pathway are enriched in both UW228-3+ and D425+ cells (Figure 5H and 5I, Supplementary Dataset 4). Furthermore, the biological process of axis elongation was enriched in neural stem cell culture conditions, suggesting transcriptional changes which assists the MB+ cells to respond to intrinsic signals responsible for developmental growth and organ positioning (Figure 5J, Supplementary Dataset 4). Overall, this suggests that neural stem cell culture conditions affect UW228-3 and D425 cells similarly functionally, although not via the exact same genes.

Nevertheless, among the shared upregulated genes, we found COL6A1, COL6A2, SEMA3A, EFNB1, ECE1, IL13RA2, TNC, and TRIB2 (Figure 5K, Supplementary Figure 5C), all of which have been associated with cancer cell stemness, migration, invasion, aggressiveness, and poor prognosis in various cancer types, including glioma [26-33]. Notably, SEMA3A functions as an axon guidance factor, playing a pivotal role in navigating the axonal network during neural development, while also regulating synaptogenesis and synaptic plasticity [34]. Importantly, autocrine SEMA3A has been demonstrated to modulate substrate adhesion, promoting glioblastoma spread, and acting as a mitogen for glioma stem cells through TGFβ-signaling activation [35, 36]. Similarly, EFNB1 promotes medulloblastoma migration and invasion and plays a crucial role in neural progenitor migration, as well as maintaining the neural progenitor cell state [37-39]. Importantly, high SEMA3A and EFNB1 expression correlate with lower overall survival in medulloblastoma patients (Figure 5L). Altogether, this shows that altering the culture conditions to neural stem cell conditions rewires the transcriptome and confers a more migratory neuronal phenotype, which may explain the increased preference for migrating to the brain area.

### Blastula-Stage Transplanted Zebrafish Embryos as a High-Throughput Orthotopic Model for In Vivo Drug testing in Medulloblastoma

Next, we evaluated the model’s utility for high-throughput drug efficacy testing. Drug testing for brain tumors faces significant challenges due to the presence of the blood-brain barrier (BBB), limiting access for many drugs. Significantly, zebrafish embryos do not establish a functional BBB until four days post-fertilization [40]. This provides a unique testing window for assessing the efficacy of drugs that might not cross the BBB, serving as a proof-of-principle platform for a compound’s effectiveness against MB cell growth. To enhance the quantitative analysis of tumor cell growth, we first transduced cells with a luciferase gene, enabling bioluminescence measurement in a plate-format. Zebrafish embryos were transplanted at the blastula-stage with either tertiary tNES-Luc or UW228-3-Luc cells. Twenty-four hours post-transplantation, embryos with cells located in the brain area were selected for drug treatment. Forty-eight hours after drug treatment, embryos were imaged and only living embryos were analyzed for bioluminescence activity (Figure 6A). The results demonstrated that the SMO inhibitor Sonidegib, as well as 4-HCP (the active metabolite of cyclophosphamide), significantly reduced the cell viability of tertiary tNES and UW228-3 cells (Figure 6B and 6C). This highlights the zebrafish MB model as a rapid and effective method for assessing drug efficacy in an in vivo setting.

**Figure 6.**
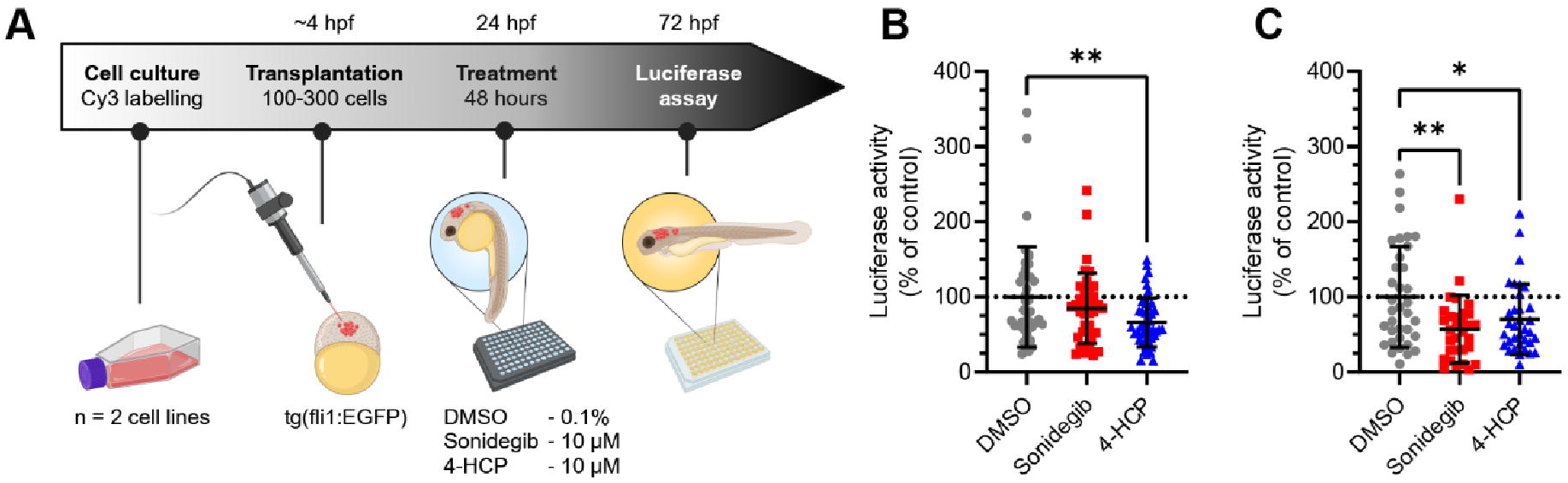
Transplantation in the blastula-stage of zebrafish embryos can be used as an orthotopic medulloblastoma model for high-throughput in vivo testing of drug compounds. **A)** Schematic overview of the treatment protocol. **B)** Luciferase activity of transplanted tertiary tNES cells **C)** and UW228-3 cells after 48 hours of treatment with sonidegib or 4-HCP. Data normalized to the DMSO control and presented as mean ± SD with each point representing one zebrafish embryo of two individual experiments. *P<0.05, **P<0.01, one-way ANOVA with Dunnett post-hoc test.

## Discussion

Here we present a zebrafish embryo model for medulloblastoma that can be used in a high-throughput format to study the kinetics of tumor cell growth, neurotropism, and drug efficacy. The injection of MB cells into blastula-stage zebrafish embryos enables a fast and efficient orthotopic transplantation of tumor cells. Importantly, the injected MB cells preferentially home to the brain and especially to the hindbrain region, the part that later develops into the cerebellum. It has previously been shown that the blastula zebrafish embryo contains homing cues that correctly guide transplanted human melanoma and glioma cells [17, 23]. We demonstrate here the importance of pre-transplantation culture conditions in enhancing the homing capacity of MB cells to the brain region. Many cell lines traditionally grow in standard culture media containing 10% fetal bovine serum (FBS). However, FBS not only exhibits batch-to-batch variability but also harbors unknown components that may impact experimental outcomes. Thus, opting for a more defined, cell type-specific media is crucial for enhancing the translatability and reproducibility of experiments. In particular, studies have shown that glioma cells cultured in serum-free media enriched with neuronal supplements B-27 and N2 as well as growth factors EGF and FGF2 show similar proliferation, migration and invasion properties to the tumors from which they originate [41-43]. Similarly, we found that the transcriptome changed significantly when we switched the growth conditions of UW228-3 and D425 to a neural stem cell-like media. Importantly, we observed upregulation of genes encoding proteins involved in cell-cell and cell-matrix contact, including integrin-1 interacting proteins. Integrins play a central role as a neurogenic factor in regulating the proliferation, stemness and migration of neural stem/progenitor cells and contribute to the formation of the neurogenic niche [44-46].

In addition to primary brain tumors, secondary brain tumors, i.e. metastases from other primary malignancies such as lung, breast, melanoma, colon and kidney cancer, occur with an incidence of 10-40% in patients with solid cancers and correlate with poor overall survival [47, 48]. The preference for metastatic colonization of the brain, neurotropism, depends on the ability of circulating tumor cells disseminated from the primary tumor site to extravasate the vasculature, cross the BBB, and adapt to the brain microenvironment while altering the extracellular matrix (ECM) and resident cells to create their own metastatic niche [49]. Interestingly, many of the commonly upregulated genes in UW228-3 and D425 cells cultured in neural stem cell media, including IL13RA2, COL6A1, and COL6A2, have been shown to promote or correlate with breast cancer metastasis to the brain [26, 27], suggesting that neural stem cell growth condition promote neurotropism. In line with this, we found that neural stem cell culture conditions resulted in the upregulation of guidance molecules SEMA3a and EFNB1, and that high expression of SEMA3A and EFNB1 in MB patients correlated with a poor prognosis. Interestingly, blocking antibodies against SEMA3A have been shown to inhibit glioblastoma progression in a mouse model [50], suggesting that it could be an interesting target for MB to further explore.

In summary, we demonstrate that early blastula-stage zebrafish embryos are a valuable orthotopic model system for medulloblastoma. This model is useful not only for assessing drug responses but also to investigate neurotropism mechanisms associated with both primary brain tumors and brain metastasis. Importantly, we show that patient-derived cells can successfully graft and grow in the zebrafish model, thus allowing for rapid expansion and testing of primary patient samples. This opens opportunities for personalized drug sensitivity screens, allowing rapid testing that can provide tailored treatment strategies for individual patients.

## Funding

Cancerfonden (22 2236 Pj), Barncancerfonden (PR2021-0080), Radiumhemmets Forskningsfonder (#214173), Vetenskapsrådet (2020-1427), and Karolinska Institutet (2-5586/2017, 2-1060/2018).

## Conflict of interest

The authors declare no conflict of interest.

## Authorship

N.v.B. and M.W. designed the study; N.v.B., A.S.O., S.L., L.Bo., L.Z., and L.Br. performed research; N.v.B. and A.S.O. analyzed data; M.W., L.Br., J.I.J. contributed new reagents; N.v.B. and M.W. wrote the original draft of the manuscript. All authors have reviewed, read, and approved the submitted version of the manuscript.

## Data availability

All data generated or analyzed during this study are included in this published article (and its Supplementary Information files) or deposited at GEO database, accession number pending.

## Acknowledgements

Figures 1A, 5A, 6A, and Supplementary Figure 1A were created with BioRender.com (https://biorender.com). We would like to thank the zebrafish core facility at Karolinska Institutet for their technical expertise and assistance and everyone in the Wilhelm lab for helpful discussions.

## Notes

### Competing Interest Statement

The authors have declared no competing interest.

