## Supplemental Figures for "Development of an orthotopic medulloblastoma zebrafish model for rapid drug testing"

**Supplementary Figure 1**

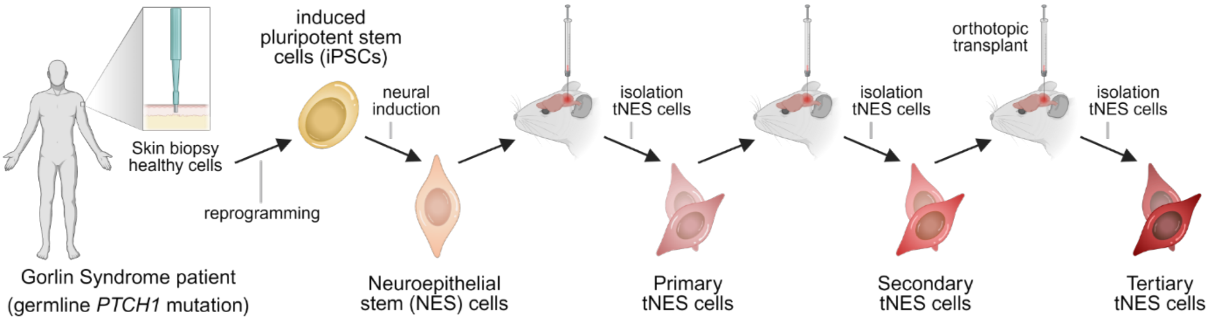

**Supplementary Figure 2**

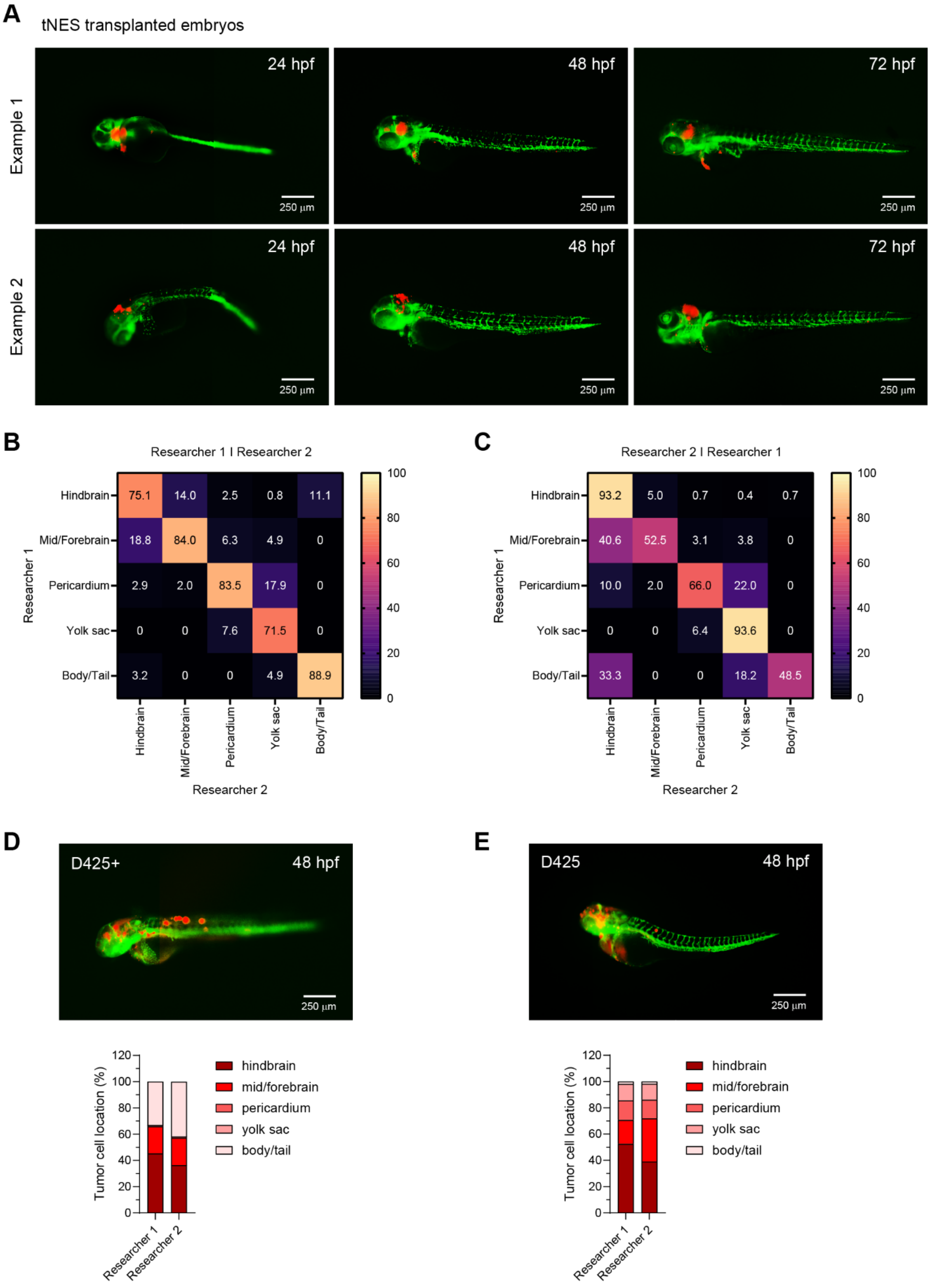

**Supplementary Figure 3**

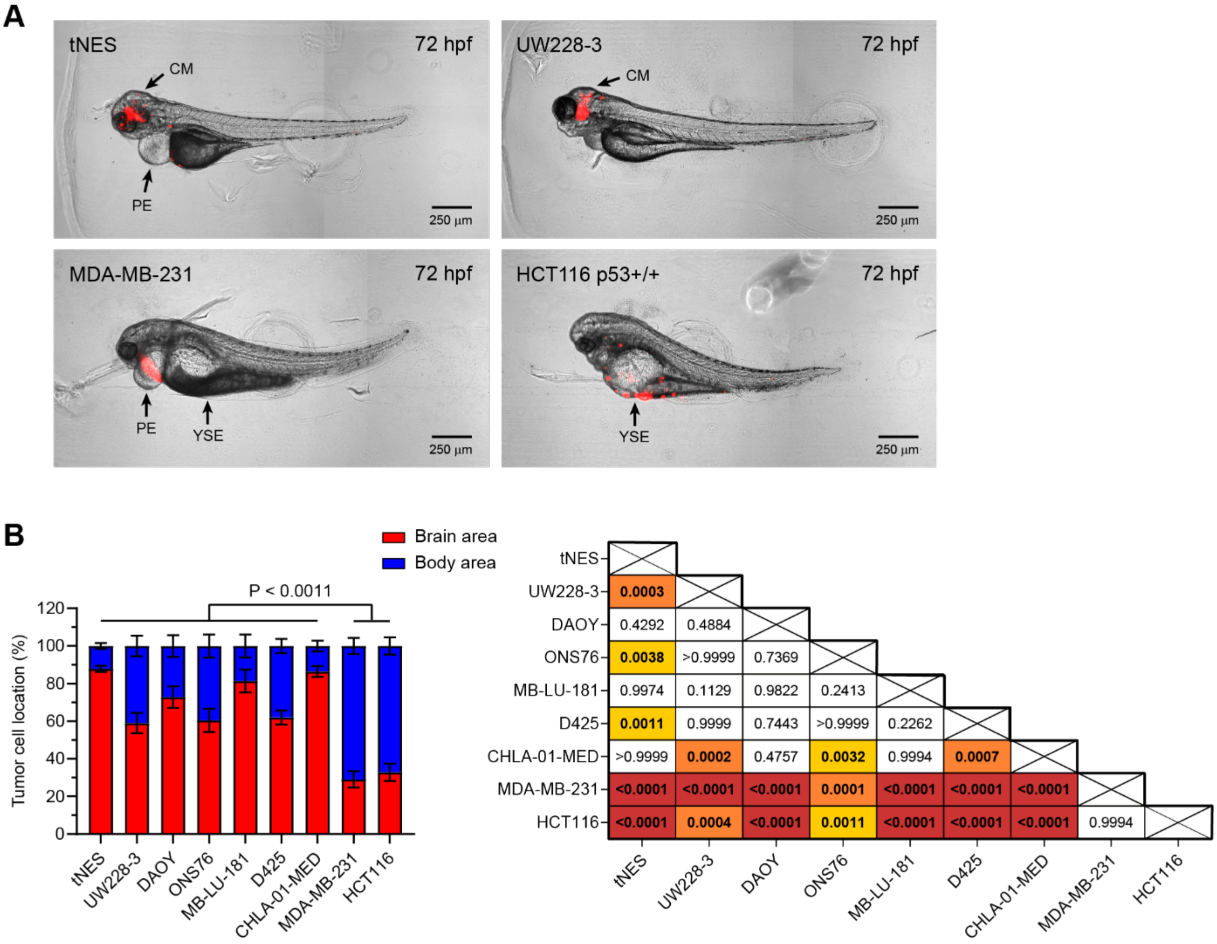

**Supplementary Figure 4**

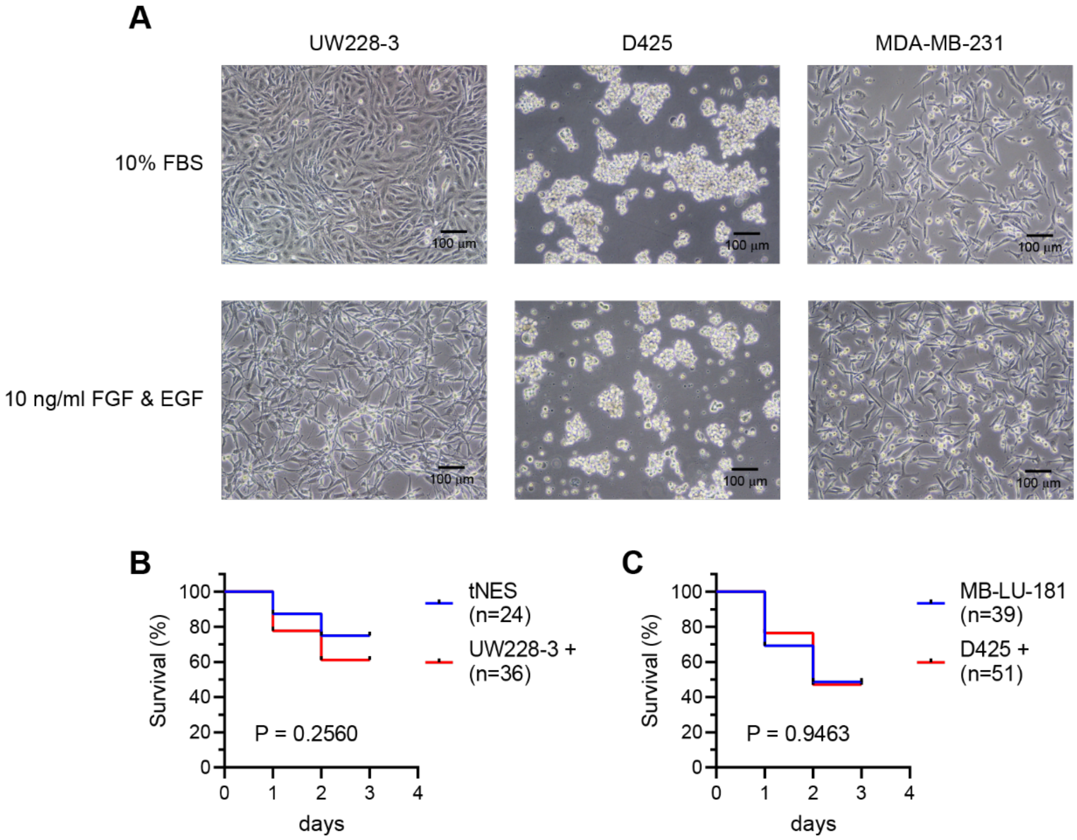

**Supplementary Figure 5**

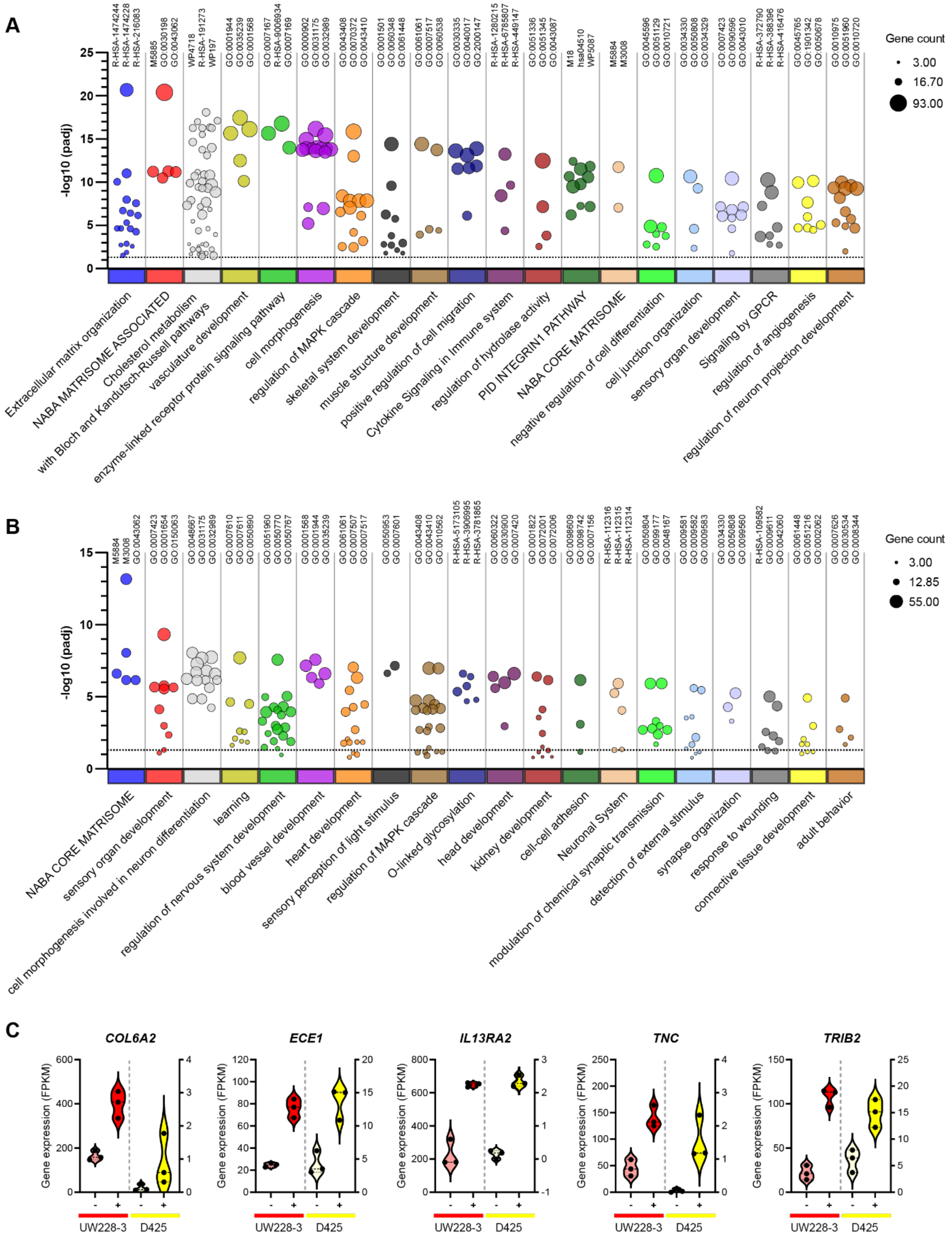

### Supplementary Figure legends

**Supplementary Figure 1. Schematic overview of the Gorlin syndrome patient-derived SHH-MB model.** Healthy skin fibroblasts were obtained through skin biopsy of the Gorlin syndrome patient carrying a germline *PTCH1* mutation. These cells were reprogrammed in induced pluripotent stem cells (iPSCs) followed by neural induction into neuroepithelial stem (NES) cells. Tumors were derived upon orthotopic transplantation of these Gorlin syndrome patient-derived NES cells, which mimic human SHH-MB on transcriptomic and phenotypical level. Re-transplantation of these tumor NES (tNES) cells resulted in increased malignancy. This model has been described by Susanto et al. [13]

**Supplementary Figure 2. Related to Figure 1. A)** Representative examples of transplanted zebrafish embryos followed over a time period of 72 hours; images were taken every 24 hours. **B)** Confusion matrices of image classification in five different areas (hindbrain, mid/forebrain, pericardium, yolk sac, body/tail) performed by researcher 1, **C)** and researcher 2. **D)** Representative image of a zebrafish with differently classified locations. Researcher 1 classified the majority of tumor cells in the hindbrain area (45.3%) whereas most tumor mass was classified in the body/tail area (41.8%) by researcher 2. **E)** Analysis of tumor cell location of a representative zebrafish image with high spread of tumor localization.

**Supplementary Figure 3. Related to Figure 2 and 3. A)** Commonly observed embryonic malformations developing upon transplantations of MB and non-MB cell lines. CM = cranial malformation, PE = pericardial edema, YSE = yolk sac edema. **B)** Comparison of brain/body distribution of transplanted MB and non-MB cell lines. one-way ANOVA with Dunnett's post-hoc test. The right panel shows the statistical results of the ANOVA test. Red = P<0.0001 (****); Orange = P<0.001 (***); Yellow = P<0.01 (**).

**Supplementary Figure 4. Related to Figure 4. A)** Representative images of UW228-3, D425, and MDA-MB-231 cells cultured in medium with 10% FBS or neural stem cell medium containing 10 ng/ml FGF and EGF. **B)** Overall survival analysis of zebrafish transplanted with UW228-3+ versus tertiary tNES cells, and **C)** D425+ versus MB-LU-181 cells. Kaplan-Meier curves depict differences in survival and statistical differences were determined using the log rank Mantel-Cox test.

**Supplementary Figure 5. Related to Figure 5. A)** Bubble plot showing the significant, upregulated gene sets belonging to each cluster. Individual clusters are separated on the x-axis and the y-axis displays the adjusted P-value. The size of the bubble indicates the overlap between the genes within the gene lists and the genes within the gene sets. The terms of the top three gene sets from each cluster are shown at the top of the plot. **B)** Bubble plot showing the significant, downregulated gene sets belonging to each cluster. **C)** Violin plots showing the DEGs *COL6A2*, *ECE1*, *IL13RA2*, *TNC*, and *TRIB2* that are significantly enriched in both UW228-3+ and D425+, expressed as fragments per kilobase (FPKM).
